# peleke-1: A Suite of Protein Language Models Fine-Tuned for Targeted Antibody Sequence Generation

**DOI:** 10.1101/2025.10.16.682644

**Authors:** Nicholas Santolla, Trey Pridgen, Prbhuv Nigam, Colby T. Ford

**Affiliations:** University of North Carolina at Charlotte, School of Data Science, Charlotte, NC, USA; University of North Carolina at Charlotte, Center for Computational Intelligence to Predict Health and Environmental Risks (CIPHER), Charlotte, NC, USA; University of North Carolina at Charlotte, Department of Bioinformatics and Genomics, Charlotte, NC, USA; North Carolina School of Science and Mathematics, Durham, NC, USA; Tuple LLC and Silico Biosciences, Charlotte, NC, USA

**Keywords:** Antibodies, Drug Design, Protein Language Model Protein Folding, Artificial Intelligence

## Abstract

The discovery of therapeutic antibodies is a traditionally arduous process. Today, the lab-based process of antibody discovery consists of several time-consuming steps that involve live animal immunization, B-cell harvesting, hybridoma creation, and then downstream engineering and evaluation. However, the use of artificial intelligence in drug design has previously been shown effective in the rapid generation of protein-specific binders, small molecules, and even antibody therapeutics, thereby replacing some of the primary steps of the drug discovery process.

Here we present *peleke-1*, a suite of protein language models fine-tuned from state-of-the-art large language models using curated antibody-antigen complex data. These models generate targeted antibody Fv sequences for a given antigen sequence input at-scale. This suite of models provides a reliable, artificial intelligence-driven approach for *in silico* therapeutic antibody discovery along with an open-source framework for future antibody language model tuning.

## Introduction

Traditional antibody discovery is slow, expensive, and resource-intensive. This typically begins with antigen preparation, followed by animal immunization to elicit an immune response. Antibody-producing B-cells are then harvested and either fused with myeloma cells to create hybridomas or incorporated into phage display libraries to capture antibody diversity.

Once sufficient antibody candidates are expressed, high-throughput screening is performed to evaluate binding affinity, specificity, and stability. Promising candidates are then subjected to engineering and optimization to improve pharmacokinetics, reduce immunogenicity, and enhance target binding. Finally, extensive *in vitro* and *in vivo* validation is required prior to clinical development for humans.

Recent advances in computational biology, particularly the rise of protein language models (PLMs), offer a new paradigm. By learning rich sequence representations from massive protein corpora, PLMs can generalize structural and functional properties of proteins, enabling predictive and generative tasks once thought infeasible. Previously, PLMs such as *ESM* (1, 2) and *ProteinMPNN* (3) have shown exceptional performance in generating realistic amino acid sequences from evolutionary-driven or structure-driven weights for general proteins.

In the antibody domain, language models such as *IgBert* (4), *AbLang* (5), *AntiBERTy* (6), and *IgGM* (7), have shown promise for accelerating candidate discovery, reducing dependence on animal immunization, and exploring immunoglobulin sequence space beyond what is accessible through natural immune repertoires or more general PLMs. However, a limitation of some of these models is that they are not trained on antibody-antigen complexes due to the lack of publicly available data.

Here we introduce *peleke-1*, a suite of protein language models fine-tuned for targeted antibody sequence generation. *peleke-1* enables the rapid design of antibody candidates conditioned on an antigen input with desired epitopes, bridging the gap between large-scale pretraining and domain-specific fine-tuning for therapeutic discovery. This suite of models provides a reliable, artificial intelligence-driven approach for *in silico* therapeutic antibody discovery along with an opensource framework and training dataset for future antibody model tuning by the computational structural biology community.

## Methods

The *peleke-1* suite consists of multiple protein-language models (PLMs), fine-tuned from existing large language models (LLMs) that span varying architectures and parameter magnitudes. To perform the fine tuning, copious antibody-antigen sequence information was collected to form a curated training dataset. The overall workflow is shown in Figure 1.

**Fig. 1.**
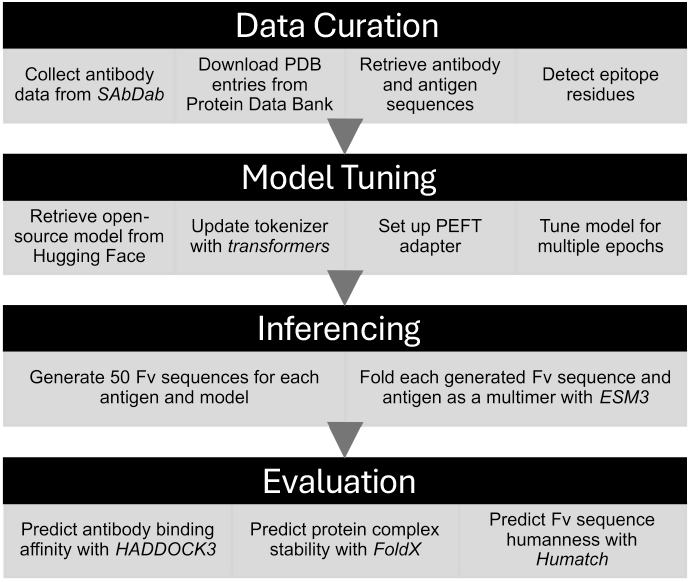
Model tuning and evaluation workflow.

### Data Curation

Antibody-antigen complexes were collected from the Structural Antibody Database (SAbDab). The SAbDab data includes the PDB ID for the complex structure hosted on the Protein Data Bank website along with heavy and light chain and antigen chain identifiers. Also, this data includes other metadata about each antibody. This dataset included 18,971 antibodies across 16,263 target antigens.

For each record, the PDB ID was used to retrieve the structure and sequence information from Protein Data Bank, filtering out any records where any of the antibody or antigen chains are missing. This resulted in a curated dataset of 9,523 complete entries that consisted of a heavy chain Fv sequence, light chain Fv sequence, and a target antigen sequence for each antibody. Example derived values are shown in Supplementary Table 2.

### Active Residue Determination

Once the collection of antibody-antigen sequences was curated, the representative PDB structures were analyzed using PandaProt (8), which identifies epitope residues on the antigen that formed polar contacts with a CDR loop residue on the Fv structure. These interfacing residue numbers were collected and used to annotate the input sequences prompt, surrounding the interfacing residues with square brackets (“[]”). Prompt Example: `Antigen: …FS[S][F][V]L[N]WY…\nAntibody: QVQL…|DIQM…`

### Fine-Tuning

Using the *transformers* package from Hugging Face, three LLMs were fine-tuned to the curated training dataset as listed in Table 1.

**Table 1.** Percentage of generated antibodies that are predicted to be human per Humatch.

| Base Model | Source | Params | PEFT Modules | Params Tuned | Trainable % |
| --- | --- | --- | --- | --- | --- |
| Phi-4 | Microsoft (9) | 14.7B | o/qkv_proj | 7,372,800 | 5.03% |
| Llama-3.1-8B-Instruct | Meta (10) | 8.03B | o/qk/v_proj | 2,097,152 | 2.61% |
| Mistral-7B-Instruct-v0.2 | Mistral (11) | 7.24B | o/qkv_proj | 13,631,488 | 0.19% |

First, the tokenizer for each model was updated to handle standard 20 amino acid characters and the Fv-chain delimiter “|”. Special tokens were added for denoting the epitope residues (`<epi>` and `</epi>`). The token embeddings of the model were then resized to accommodate these new tokens.

Then, the training data are formatted and tokenized using the updated tokenizer. For example, `KSF[E][D]AKCAA…` becomes `K,SF,<epi>, E,</epi>,<epi>,D,</epi>,AK,CAA,…` after tokenization through the updated *Phi-4*-based tokenizer. Stop tokens were also used at the end of the antigen sequence and antibody sequences to guide the LLM’s response.

*Phi-4* and *Llama-3*.*1-8B-Instruct* were tuned on a cloud-based NVIDIA H100 GPU in Microsoft Azure Machine Learning and *Mistral-7B-Instruct-v0*.*2* was tuned on NVIDIA 5090 and 3090ti dual GPUs locally.

Each LLM was fine-tuned to minimize loss over a few epochs, stopping when stable Fv-like sequences were consistently being generated for each chunk of steps. The models were then checkpointed for future use, evaluation, and inferencing.

### Evaluation

Upon sufficient tuning of the PLMs, a test set of antigen sequences was selected as benchmarks for the antibody generation. The target benchmark antigens used were:

#### Oncotargets

- EGFR: Human epidermal growth factor receptor. (PDB: 8HGO)
- PD-L1: Programmed death-ligand 1, an immune checkpoint inhibitor. (PDB: 4Z18)
- PD-1: Programmed death protein 1, an immune check-point protein. (PDB: 5JXE)

#### Infectious Disease Proteins

- MBP: Maltose-binding protein from *E. coli*, a common protein expression tag. (PDB: 1NL5)
- BHRF1: BCL-2 homolog from Epstein-Barr virus, an anti-apoptotic protein. (PDB: 2WH6)

#### Others

- IL-7Rα: Interleukin-7 receptor alpha chain, a cytokine receptor subunit. (PDB: 3DI3)
- BBF-14: A synthetic 112-AA β-barrel protein. (PDB: 9HAC)

This set encompasses 6 standard benchmark antigens to match AdapytvBio’s BenchBB service (BHRF1, EGFR, IL-7Ra, MBP, PD-L1, and BBF-14) (12) and 1 additional antigen (PD-1).

For each antigen, 50 novel Fv antibody sequences were generated. Each paired set of Fv sequences were co-folded with their respective antigen using the *esm3-medium-multimer-2024-09* model from Evolutionary Scale (2).

Each Fv sequence was also quality-checked using multiple methods. This included a test to number the sequences through the Chothia numbering system using the ANARCI library (13) (to ensure Fv-like sequences), an analysis for evaluating humanness with Humatch (14), and stability predictions with FoldX (15). This closely follows the evaluation logic as published in Santolla and Ford, 2025.

HADDOCK3’s *haddock3-score* method was used to predict the binding affinity of the predicted Fv structure to the target antigen (17, 18). Then, these affinity metrics were collected to assess the generated Fv structure’s binding to the desired antigen. The metrics are as follows:

- Van der Waals intermolecular energy (*vdw*) in kcal/mol
- Electrostatic intermolecular energy (*elec*) in kcal/mol
- Desolvation energy (*desolv*) in kcal/mol
- Buried surface area (*bsa*) in Å^2^
- Total energy (*total*): 1.0*vdw* + 1.0*elec* in kcal/mol
- HADDOCK score: 1.0*vdw* + 0.2*elec* + 1.0*desolv* + 0.1*air*

Evaluation outputs were then summarized by base LLM and antigen to assess overall model performance.

## Results

Across the 7 benchmark antigens, 50 Fv sequences were generated for each of the 3 fine-tuned models, a total of 1,050 test complexes. The tuned models were inferenced locally on NVIDIA 5090 and 3090ti GPUs. Inferencing took <1 minute per Fv generation. Note that each model was inferenced 350 times (7 benchmark antigens*×*50 runs) to generate pairs of heavy and light chain Fv sequences in each run.

### Antibody Realness and Stability

Each model consistently produced Fv heavy and light chain sequences, 100% of which were successfully numbered using the Chothia numbering scheme (with only 4 eliciting duplicate residue numbering warnings).

Furthermore, the models consistently generate human-like sequences, with 82% and 80% of Fv sequences predicted to be human for heavy and light chains, respectively. See Supplementary Table 1 for a full breakdown by model and antigen.

Regarding protein stability, the generated antibody-antigen complexes averaged a total energy of −215.59 kcal/mol, polar solvation of −715.03 kcal/mol, and hydrophic solvation of 1,246.67 kcal/- mol across all models and antigens. These indicate predicted stable binding and solubility. While overall stability was similar across models, antigen-level metrics varied. EGFR had the most stable interactions and BBF-14 had the least. See Supplementary Figure 1 for the stability metric distributions by model and antigen.

### Sequence Diversity

Due to the nature of language models, output sequence lengths varied. The generated heavy chain sequences were between 103 and 145 amino acids in length 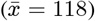 and light chains were between 98 and 116 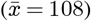 amino acids.

Conversely to our previous work in Santolla and Ford, 2025 that used a Microsoft EvoDiff diffusion model (19), where the conditionally-diffused sequences exhibited relatively low sequence diversity, the *peleke-1* language models generated sequences with significant diversity. In the generated heavy chain sequences, various standard antibody patterns were common (e.g. sequences starting with EVQL or QVQL and ending with TVSS), though there was significant diversity in the other regions. In the generated light chains, lower diversity was experienced outside the CDR loop regions with standard DIQM or DIVL beginning patterns and ending with TKVEIK. See Supplementary Figure 2 for full WebLogos.

### Binding Performance

As shown in Table 2, the models were better overall at generating antibodies for some target antigens compared to others. For example, antibodies against targets MBP, PD-1, and PD-L1 had much better (lower) affinity scores and van der Waals energies as compared to those generated for BBF-14, BHRF1, and EGFR.

**Table 2.** Curated antibody-antigen complex training data from SAbDab.

| Antigen | *peleke-1* Fine-Tuned Model |  |  |
| --- | --- | --- | --- |
|  | *llama-3.1-8b-instruct* | *mistral-7b-instruct-v0.2* | *phi-4* |
| BBF-14 | 22.73% (63.34) | 15% (106.84) | 18.42% (49.88) |
| BHRF1 | 2.63% (218.86) | 10.81% (289.61) | 9.3% (131.62) |
| EGFR | 20% (101.64) | 23.26% (24.66) | 20% (163.88) |
| IL-7Ra | 36.17% (-3.46) | 46.81% (-10.11) | 31.91% (13.64) |
| MBP | 67.39% (-52.56) | 57.78% (-52.52) | 69.39% (-57.98) |
| PD-1 | 73.47% (-66.72) | 79.59% (-82.51) | 76.6% (-70.12) |
| PD-L1 | 79.59% (-71.53) | 70.83% (-62.52) | 72.92% (-73.38) |

The fine-tuned models also performed similarly overall for a given antigen. As shown in Figure 2, PD-1 and PD-L1 had the best overall distributions of values, indicating stable performance of the antibody generation for those antigens specifically. Conversely, in BBF-14 and BHRF1, have the largest disparity in overall generated antibody binding.

**Fig. 2.**
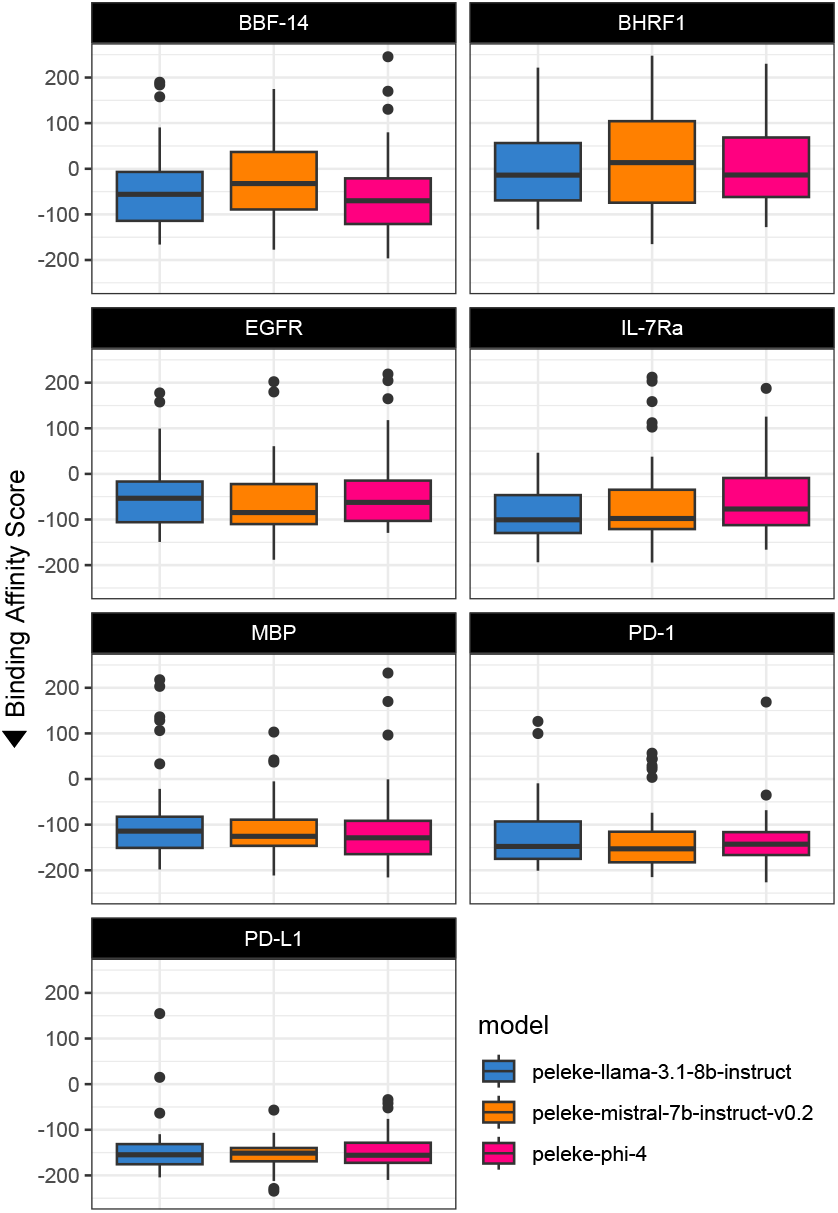
Boxplots depicting the predicted binding affinity score distribution by antigen and model. More negative values indicate better overall binding. Note that the y-axis is showing scores between [-250, 250].

### Structural Examples

Across the 1,050 generated sequences in this study, many have epitopes in the resulting predicted structures that closely match the desired epitope residues defined in the model input prompt.

For example, in the BHRF1 antigen, a normal target protein is the BCL-2-like protein, shown in yellow in Figure 3B. This interaction enables apoptosis inhibition by the Epstein-Barr virus (20). Using the *peleke-phi-4* model with an input prompt highlighting the active residues between BHRF1 and the BCL-2-like protein from PDB 2WH6, we are able to generate the antibody shown in pink in Figure 3A, which is predicted to block the interaction between BHRF1 and the BCL-2 homolog protein.

**Fig. 3.**
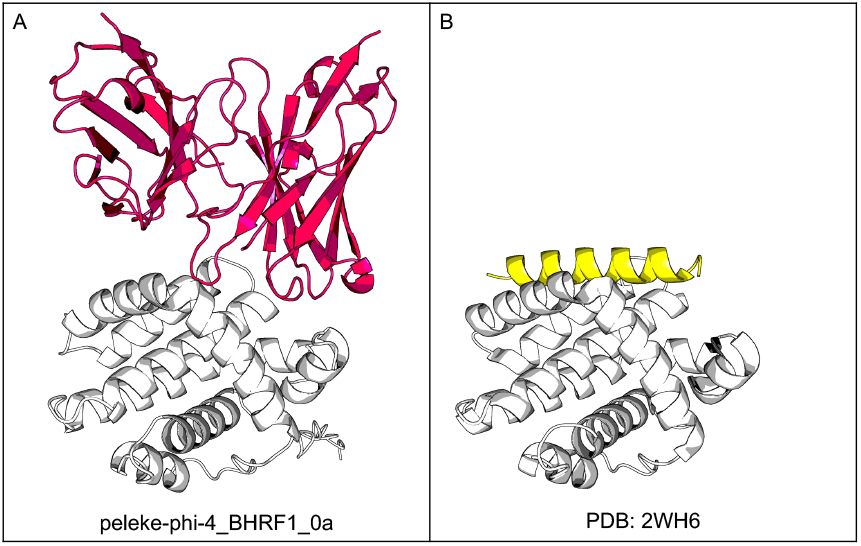
Comparison of (A) *peleke-phi-4_BHRF1_0a*, a novel generated anti-BHRF1 Fv structure, in pink; and (B) a BCL-2-like protein 11 (in yellow) bound to BHRF1 (in white, PDB: 2WH6).

One of the best predicted binders across the set of 1,050 generated sequences was *peleke-phi-4_MBP_20* with an overall binding score of −215.63 and a van der Waals energy of −102.422 kcal/mol. The antigen prompt for MBP (maltose binding protein) highlighted the residues surrounding the maltose ligand with the goal of blocking the binding pocket with the antibody. As shown in Figure 4A, this generated candidate is predicted to be a strong binder on the opposite side of the protein, missing the target epitope.

**Fig. 4.**
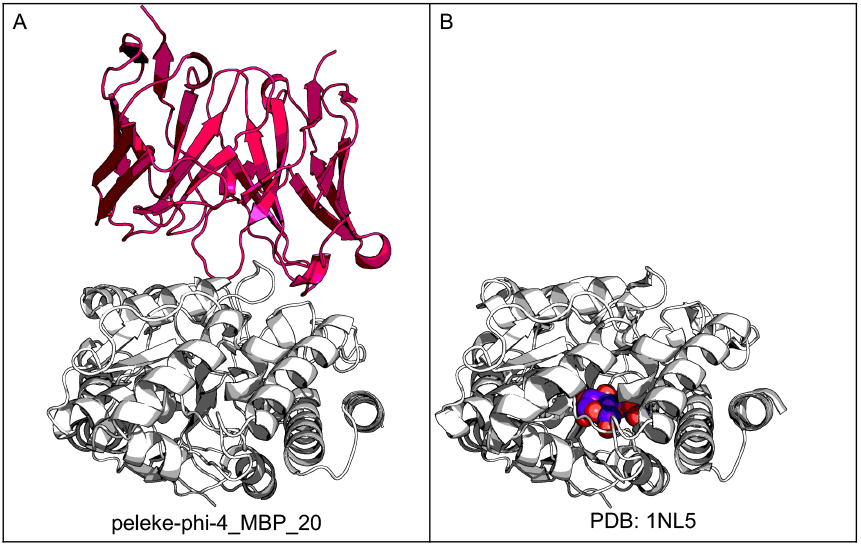
Comparison of (A) a novel anti-MBP candidate *peleke-phi-4_MBP_20* (in pink); and (B) the maltose binding protein in its closed conformation (in white) with maltose bound inside (in purple, PDB: 1NL5).

Interestingly, the side where the antibody bound is a conformationally-important site known as the “balancing interface” that controls the ligand-binding cleft to allow/block the maltose ligand (21). Thus, this novel antibody, shown in pink in Figure 4A, could play a role in inhibiting this conformational change.

Lastly, PD-1, a checkpoint inhibitor for which there are multiple approved antibody therapeutics, is a common oncotarget in various cancers. Ranging from metastatic melanoma to metastatic nonsmall cell lung cancer (NSCLC) to Hodgkin lymphoma, anti-PD-1 antibodies are used as a combination therapy with other chemotherapy drugs.

As previously reported in Ford, 2024, pembrolizumab (a leading approved anti-PD-1 antibody, marketed as Keytruda^®^ by Merck & Co., Inc.) had a predicted van der Waals energy of −62.22 kcal/mol (22). Of the 150 candidates generated by the *peleke-1* models, 83 (55.3%) of the candidates have better affinity scores than pembrolizumab and other anti-PD-1 antibodies. Shown in Figure 5 are the best candidates per model, all with predicted van der Waals energies <-100 kcal/mol.

**Fig. 5.**
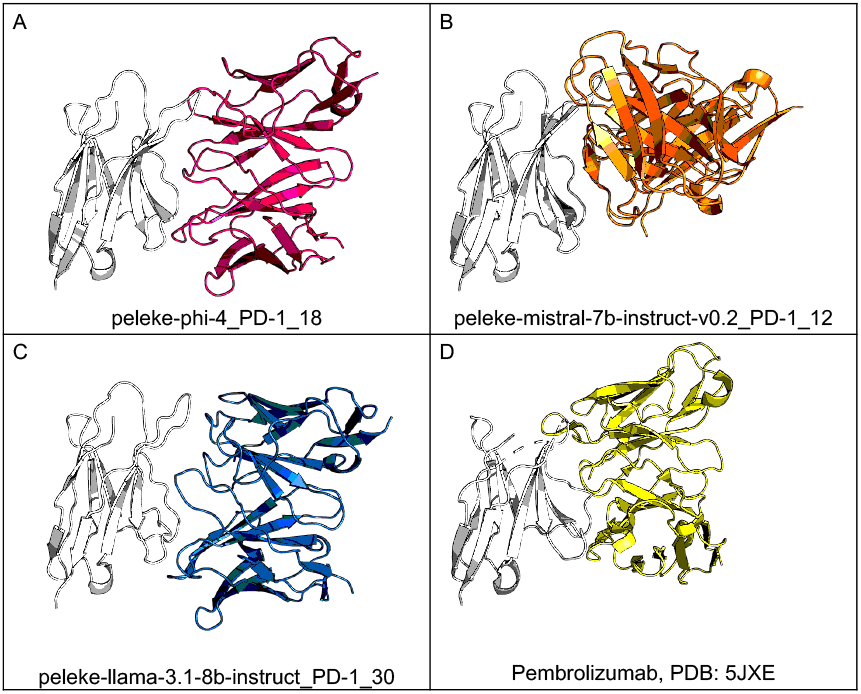
Comparison of 3 novel anti-PD-1 candidates (A) *peleke-phi-4_PD-1_18* (in pink); (B) *peleke-mistral-7b-instruct-v0*.*2_PD-1_12* (in orange); and (C) *pelekellama-3*.*1-8b-instruct_PD-1_30* (in blue); with (D) pembrolizumab (in yellow) bound to PD-1 (in white, PDB: 5JXE).

As shown in these aforementioned examples, across various targets, the *peleke-1* series of models often generates novel, targeted antibodies with strong binding potential. While additional testing across more target antigens is needed, the set of 7 benchmark antigens shown here proved useful in highlighting the learned behavior of the tuned models and where future improvements may increase performance, binding affinity, and epitope specificity.

## Discussion

The use of AI in antibody discovery poses a great benefit to the therapeutic development process. As shown in this study, specialized PLMs such as *peleke-1* provide a cost-effective method to computationally generate novel antibody sequences that reliably fold into desired structures and bind to the targets of interest. By fine-tuning pretrained language models on antibody-specific corpora, *peleke-1* bridges the gap between large-scale protein representation learning and targeted therapeutic design.

Compared to traditional workflows, which rely on animal immunization and hybridoma production, *peleke-1* can significantly reduce the experimental lab workload. In contrast to prior PLMs that have been applied broadly across proteins, our suite of models explicitly encodes antibody sequence and antigen biases, enabling more targeted generation. This domain specialization highlights the value of task-specific fine-tuning for biologics development.

Nevertheless, several limitations remain. As always, computational generation does not eliminate the need for experimental validation. Thus, candidate antibodies must still undergo biochemical characterization and functional assays. Second, while *peleke-1* demonstrates strong performance in generating binders to predefined epitopes, generalization across diverse antigen classes remains an open challenge. Finally, like other deep learning systems, model interpretability and robustness require further investigation.

The *peleke-1* project was designed for ease-of-use and flexibility for antibody discovery. By utilizing the *transformers* framework from Hugging Face, we have enabled easy inferencing for users and simplified the tuning process for future developers wanting to extend or adapt this work.

Looking forward, integrating *peleke-1* with high-throughput screening pipelines and other structure-prediction frameworks could accelerate end-to-end antibody discovery. Moreover, additional tuning time and automated evaluation, along with incorporating feedback from experimental assays into iterative model design, may further improve binding specificity and developability. Ultimately, by combining computational generation with empirical validation, approaches like *peleke-1* have the potential to transform antibody discovery into a faster, more scalable and accessible process with a higher likelihood of therapeutic success.

## Key Contributions

This work showcases the following contributions:

1. Domain-specialized PLMs for antibodies: We introduce *peleke-1*, a suite of protein language models fine-tuned on antibody-specific corpora to enable targeted sequence generation.
2. Epitope-conditioned design: We demonstrate that *peleke-1* can generate antibody sequences conditioned on desired epitopes, enabling controllable design beyond general-purpose protein models.
3. Bridging AI and molecular generation: By treating antibodies as structured sequences of amino acid tokens, we highlight how task-specific fine-tuning of PLMs will accelerate therapeutic discovery.
4. Open-sourced for the community: We release model weights, curated training data, tuning code, and generation pipelines to promote reproducibility and enable further research in machine learning for antibody engineering.

## Contributors

Authors NS and CTF developed the *peleke* fine-tuning pipeline. TP and CTF developed the training data preparation logic and evaluation pipeline. PN assisted in the research into candidate LLMs. CTF performed the evaluation of the models. All authors wrote and reviewed this manuscript.

## Acknowledgments

We acknowledge the following entities at the University of North Carolina at Charlotte: the Center for Computational Intelligence to Predict Health and Environmental Risks (CIPHER), the Department of Bioinformatics and Genomics, and the School of Data Science.

We would like to think Rafael Jaimes III from MIT Lincoln Lab for his help in reviewing this manuscript.

## Data Sharing Statement

All code, data, results, and additional analyses are openly available on GitHub at: https://github.com/silicobio/peleke. This repository includes the open-source logic for tuning additional *peleke* models.

Model weights are hosted on Hugging Face at https://huggingface.co/collections/silicobio/peleke-1-686c7e91c90f7f65a748698d. This collection of models includes the updated tokenizers and safetensors for each of the adapter models tuned and sample inferencing code.

## Supplementary Materials

**Fig. 1.**
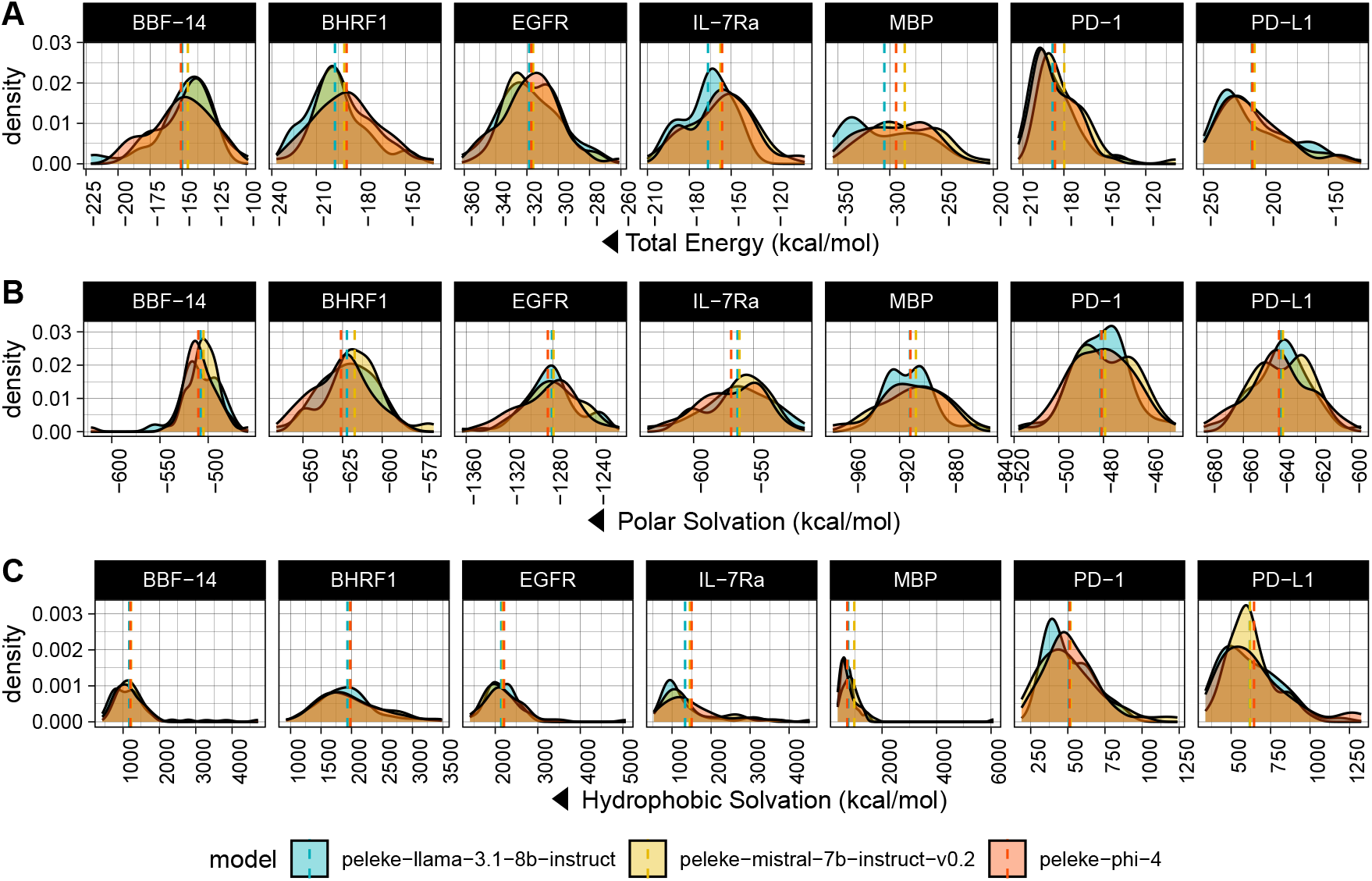
Density plots of (A) total energy, (B) polar solvation, and (C) hydrophobic solvation. These stability metrics are shown by model and benchmark antigen.

**Fig. 2.**
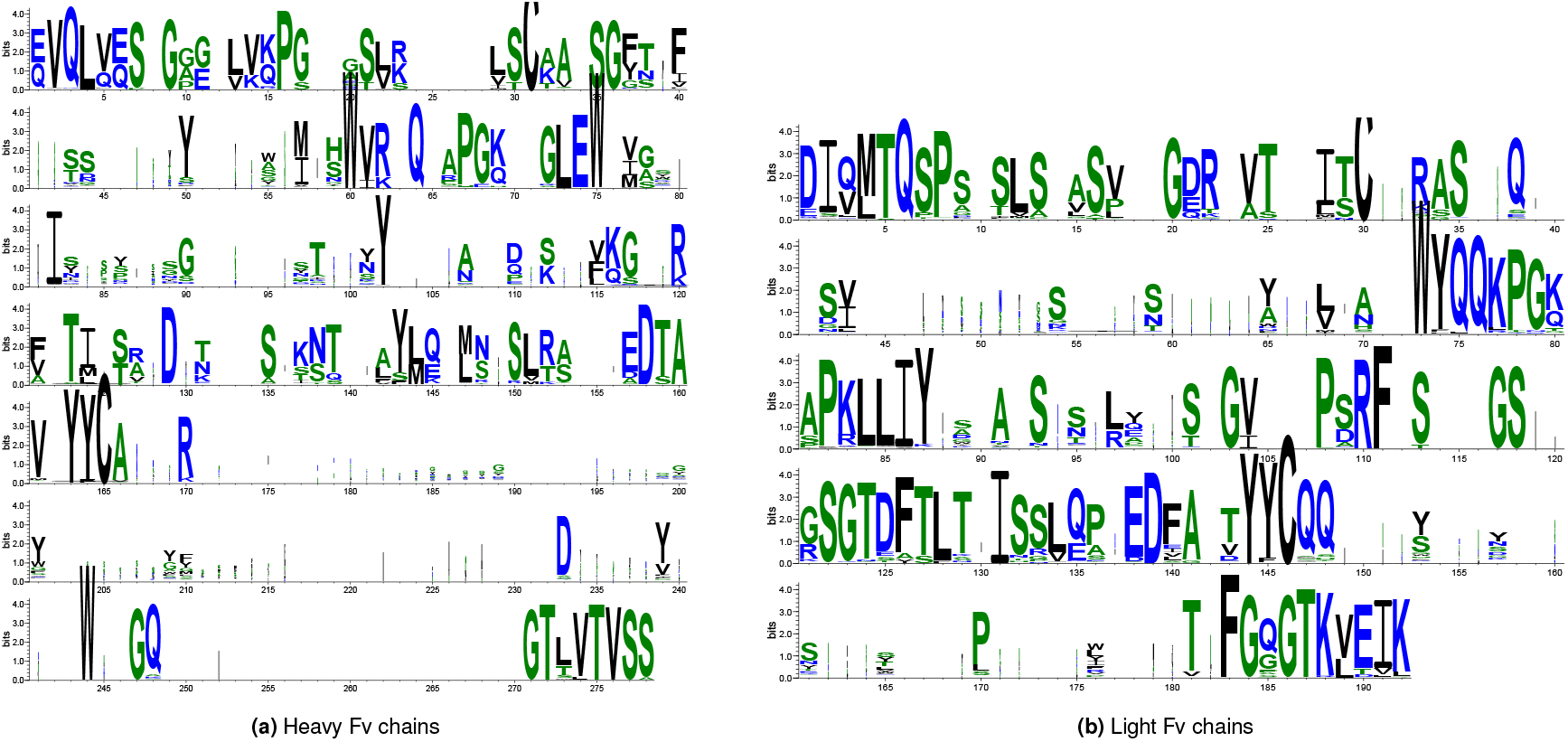
WebLogos of the generated Fv sequences (23).

**Table 1.** Base models and LoRA configurations used in this study.

| Model | peleke-llama-3.1-8b-instruct |  |  | peleke-mistral-7b-instruct-v0.2 |  |  | peleke-phi-4 |  |  | Overall |  |  |
| --- | --- | --- | --- | --- | --- | --- | --- | --- | --- | --- | --- | --- |
| Antigen | H Chain | L Chain | Paired | H Chain | L Chain | Paired | H Chain | L Chain | Paired | H Chain | L Chain | Paired |
| BBF-14 | 96% | 90% | 50% | 76% | 74% | 52% | 84% | 80% | 54% | 85% | 81% | 52% |
| BHRF1 | 88% | 88% | 62% | 70% | 76% | 46% | 80% | 78% | 54% | 79% | 81% | 54% |
| EGFR | 78% | 82% | 64% | 80% | 82% | 54% | 80% | 70% | 50% | 79% | 78% | 56% |
| IL-7Ra | 88% | 86% | 56% | 82% | 78% | 52% | 82% | 70% | 42% | 84% | 78% | 50% |
| MBP | 82% | 86% | 58% | 74% | 68% | 38% | 72% | 76% | 42% | 76% | 77% | 46% |
| PD-1 | 86% | 96% | 56% | 74% | 68% | 44% | 90% | 72% | 50% | 83% | 79% | 50% |
| PD-L1 | 86% | 86% | 58% | 86% | 90% | 58% | 84% | 88% | 56% | 85% | 88% | 57% |
| Overall | 86% | 88% | 58% | 77% | 77% | 49% | 82% | 76% | 50% | 82% | 80% | 52% |

**Table 2.**
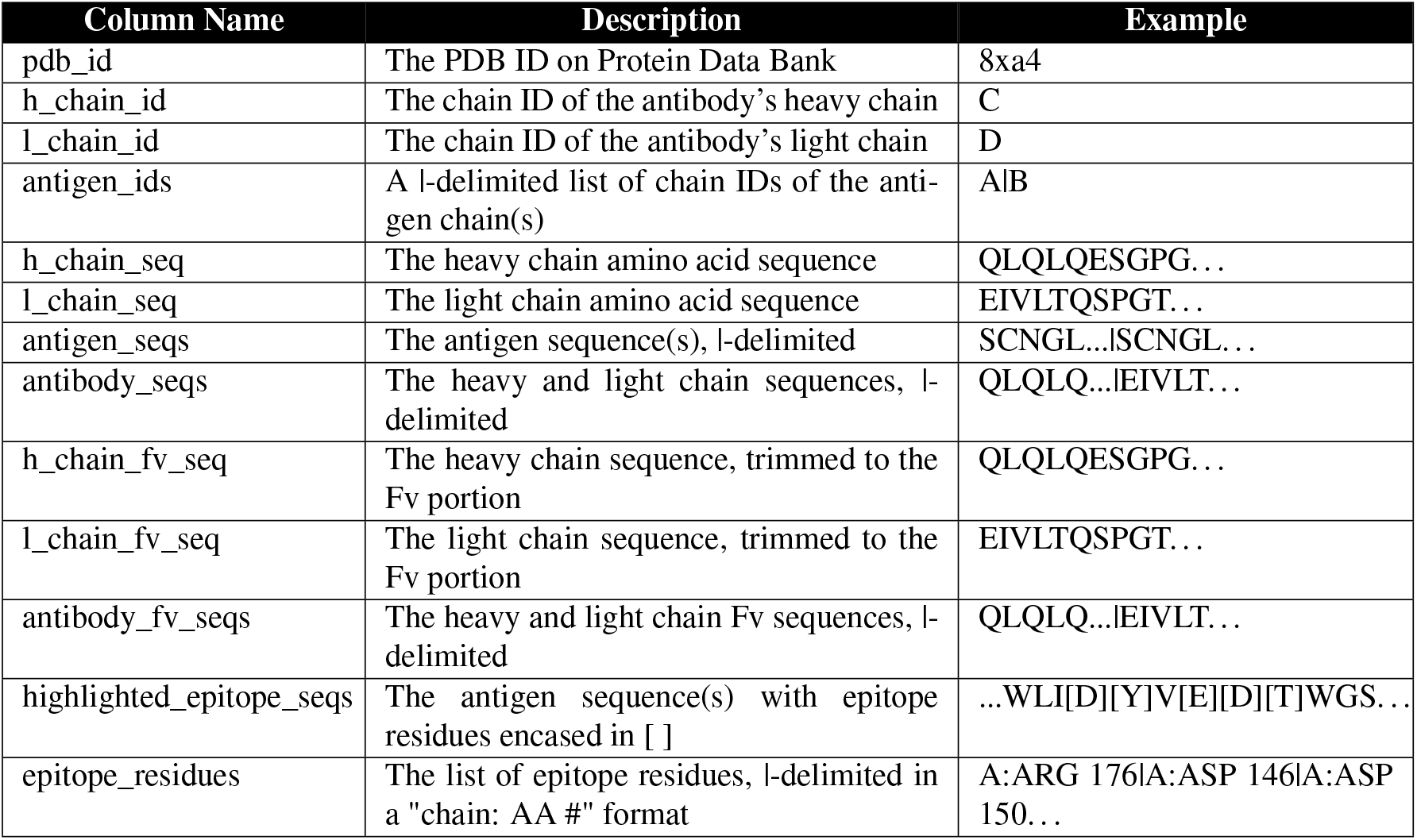
Percentage of complexes with van der Waals energies <-25 kcal/mol (median value shown in parentheses) by fine-tuned model and antigen.

